# A Comparison of Embedding Aggregation Strategies in Drug-Target Interaction Prediction

**DOI:** 10.1101/2023.09.25.559265

**Authors:** Dimitrios Iliadis, Bernard De Baets, Tapio Pahikkala, Willem Waegeman

**Affiliations:** Department of Data Analysis and Mathematical Modelling, Ghent University, Coupure links 653, Ghent, B-9000, Belgium; Department of Computing, University of Turku, Turku, 20500, Finland, Country

**Keywords:** Drug-target interaction prediction, Binding affinity prediction, Recommender systems, Deep learning

## Abstract

The prediction of interactions between novel drugs and biological targets is a vital step in the early stage of the drug discovery pipeline. Many deep learning approaches have been proposed over the last decade, with a substantial fraction of them sharing the same underlying two-branch architecture. Their distinction is limited to the use of different types of feature representations and branches (multi-layer perceptrons, convolutional neural networks, graph neural networks and transformers). In contrast, the strategy used to combine the outputs (embeddings) of the branches has remained mostly the same. The same general architecture has also been used extensively in the area of recommender systems, where the choice of an aggregation strategy is still an open question. In this work, we investigate the effectiveness of three different embedding aggregation strategies in the area of drug-target interaction (DTI) prediction. We formally define these strategies and prove their universal approximator capabilities. We then present experiments that compare the different strategies on benchmark datasets from the area of DTI prediction, showcasing conditions under which specific strategies could be the obvious choice.

## 1 Introduction

Drug discovery is a challenging task that has consistently proven to be time-consuming and expensive [1]. As a result, pharmaceutical companies are turning towards computational methods capable of automating more stages of their drug discovery pipelines. One of the earliest stages involves the prediction of interactions between chemical compounds and biological targets, also known as drug-target interaction (DTI) prediction, replacing large-scale biological screening experiments with more efficient approaches. In this area, two groups of computational methods are distinguished: docking methods and machine learning approaches. Docking methods simulate the physical interaction of molecules and proteins in the 3D space accounting for the physical structure [2]. Machine learning approaches, on the other hand, predict drug-target interactions by learning from data, using databases that contain results of traditional high-throughput screening experiments [3].

Recently, deep learning methods have gained a lot of interest in DTI prediction. In these methods, explicit features for the compounds and proteins are available beforehand and passed to a deep neural network. Even though many approaches exist, we will focus on a quite popular sub-category of two-branch deep neural networks. As shown in Fig. 1, these architectures consist of two neural network components that encode the compound and protein features, respectively. The two embedding vectors generated by the branches are then aggregated to obtain the final prediction. A nonexhaustive list of recent approaches that consider an architecture of that kind can be found in Table 1.

**Table 1:** two-branch methods.

| Method | Compound representation | Compound branch | Protein representation | Protein branch |
| --- | --- | --- | --- | --- |
| DeepConv-DTI [4] | Morgan | MLP | amino acid sequence | CNN |
| DeepDTA [5] | SMILES | CNN | amino acid sequence | CNN |
| Shin et al. [6] | SMILES | transformer | Fasta | CNN |
| MDeePred [7] | ECFP<sub>4</sub> | MLP | multi-channel protein features | CNN |
| Chen et al. [8] | molecular graph | GNN | amino acid sequence | BERT |
| Torng et al. [9] | molecular graph | GCNN | protein graph | GCNN |
| SSnet [10] | fingerprint | MLP | backbone curvature, torsion | CNN |
| Tsubaki et al. [11] | molecular graph | GNN | amino acid sequence | CNN |
| Kang et al. [12] | SMILES | BERT | amino acid sequence | BERT |
| Nguyen et al. [13] | molecular graph | GCN, GAT, GIN, GAT-GCN | amino acid sequence | CNN |
| Mingjian et al. [14] | molecular graph | GNN | protein graph | GNN |
| DeepPurpose [15] | various encodings | MLP, CNN, CNN-RNN... | various encodings | MLP, CNN, CNN-RNN... |

**Fig. 1:**
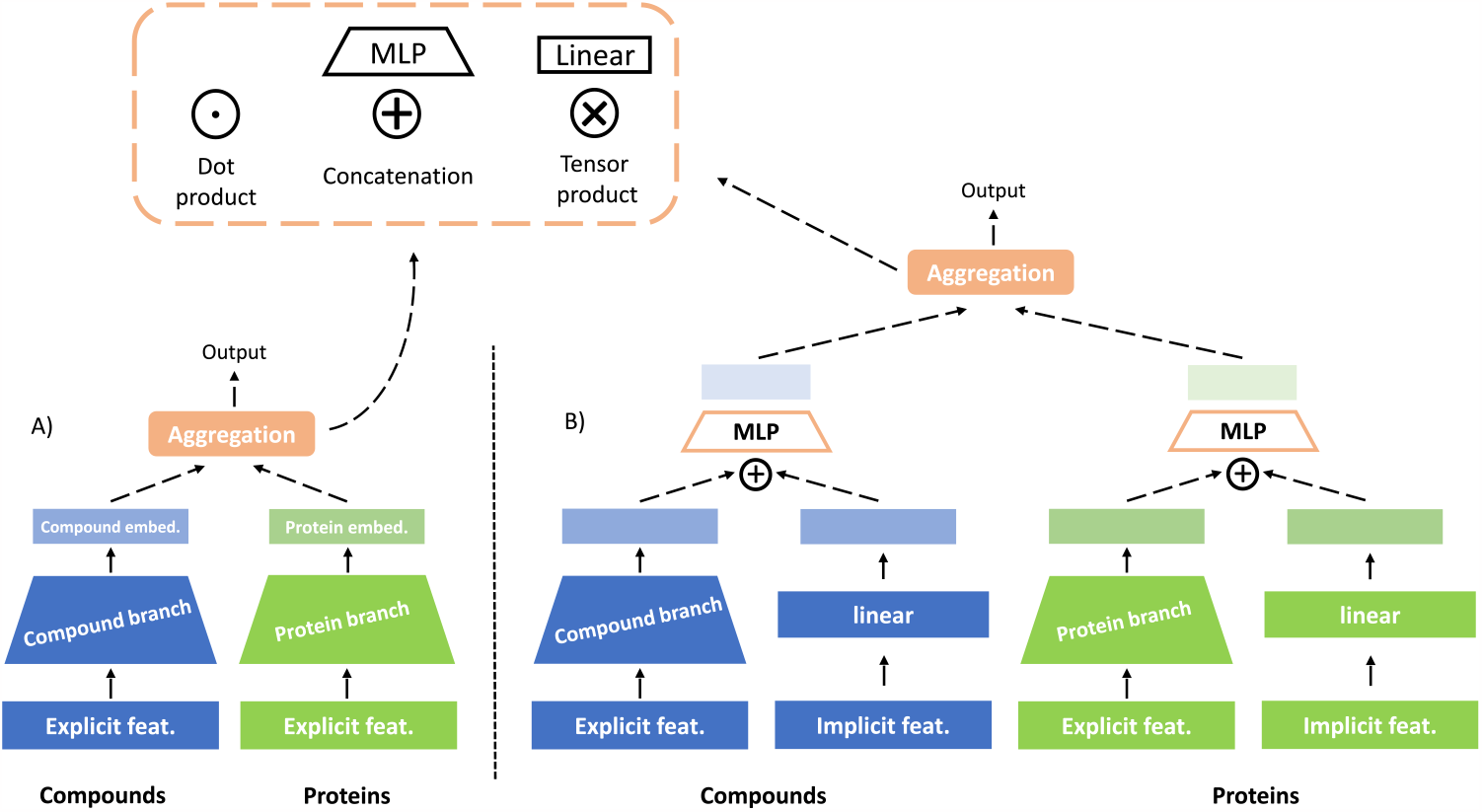
A summary of the two main versions of the architecture we used. First, the two-branch architecture (sub-figure A) encodes the explicit feature representations that are available for the compounds and proteins. The resulting embeddings can be aggregated with one of the three strategies (MLP, dot product, tensor product). The second version of the architecture combines both the implicit and explicit information available for the compounds and proteins (sub-figure B). The explicit features can be encoded in the exact same manner as shown in sub-figure A, while the one-hot encoded dummy vectors for the compounds and proteins can be transformed into dense representations using a single fully-connected layer. The intermediate embeddings from the explicit and implicit features are aggregated using the MLP strategy. This output of the MLPs is the compound and protein embeddings that can then be aggregated with any of the three available strategies.

DTI prediction is closely related to recommender systems based on collaborative filtering, such as the well-known Netflix challenge [16]. In both cases, a standard dataset takes the form of triplets:{*compound, protein, activity*} or{*user, item, interaction*} . These triplets can be arranged in a sparse matrix, and in its simplest form the prediction task is matrix completion. However, when explicit feature representations are used, three additional prediction settings become feasible, in addition to matrix completion [17]: making predictions for new compounds, new proteins or new proteincompound combinations. In DTI prediction, every method utilizes the explicit features that are available for the compounds and proteins. In contrast, similar methods in recommender systems often do not use explicit features, but create one-hot dummy vectors (implicit features) to characterize the users and items [18, 19].

DTI prediction and collaborative filtering methods also consider different aggregation strategies for the drug-target or user-item embeddings. Every DTI prediction method in Table 1 concatenates the two embeddings and then passes the resulting vector to a multi-layer perceptron (MLP strategy, see Fig. 1). Conversely, the area of collaborative filtering has a more diverse landscape, with various aggregation strategies that are regularly used, see e.g. [18, 20–26]. Especially the dot product has been extensively used in that area, and papers benchmarking the dot product versus the MLP have been published [27, 28]. Currently, a similar investigation is missing from the area of DTI prediction.

In this work, we will discuss the behavior of different embedding aggregation strategies in DTI prediction. To this end, we will analyze the above-mentioned MLP and dot product strategies, as well as a third strategy, the tensor product, which used to be popular in the era of kernel methods [29–31]. First, we will present theoretical results highlighting the universal approximation properties of all three strategies, departing from well-known mathematical building blocks. Subsequently, we will present benchmarking results of the three strategies on combinations of DTI datasets, branch types and prediction settings. Furthermore, we will investigate the effect of adding implicit feature representations, and we will interpret the learned embeddings. As a result, the main goal of this paper is to compare the aggregation strategies in detail and not to present a winning architecture that achieves state-of-the-art performance.

### 2 Methods

### 2.1 Aggregation strategies

We first describe the details of the considered aggregation strategies. As shown in Fig. 1, we compare three strategies: the dot product, the multi-layer perceptron and the tensor product.

1. **Dot product:** In the dot product strategy, the dot product of the two embeddings is directly computed and used as the prediction of the model. This strategy requires both embeddings to have the same size, a restriction that we will further investigate in a later section.
2. **Multi-layer perceptron:** The MLP strategy concatenates the embeddings and then passes the resulting vector as input to a multi-layer perceptron, which, in turn, terminates at a single output node. As stated before, the use of an MLP increases the capacity of the overall model compared to the simple dot product operation, but, perhaps, also introduces an unwanted overhead.
3. **Tensor product:** The tensor product strategy first computes the tensor product of the two embeddings and then uses a single fully-connected layer that terminates at an output node. In fact, this aggregation strategy has never been suggested as an embedding aggregation strategy for deep neural networks, but it has been extensively used in kernel methods [29–31].

For reproducibility reasons, and to describe the universal approximation capabilities of the three aggregation strategies, we present formal definitions of the three strategies. Let *𝒳* and *𝒯* be two Euclidean spaces for compounds 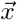 and proteins 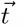, respectively. We formally define the problem of DTI prediction as that of estimating functions f : *𝒳* × *𝒯* → *𝒴*, where *𝒴* = ℝ in case of regression problems. Let us consider hypothesis spaces

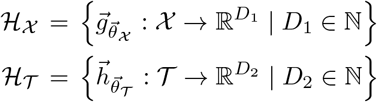

for learning compound embeddings from *𝒳* to 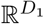 with dimensionality D_1_, and protein embeddings from 𝒯 to 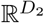 with dimensionality D_2_. 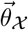 and 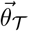 denote the parameterizations of the two types of functions. Moreover, let us consider the space C(𝒳 × 𝒯)of all continuous real-valued functions f : 𝒳 × 𝒯 → ℝ. A subspace ℌ_DP_ of C(𝒳 × 𝒯) corresponds to the dot product strategy of the two-branch architecture, *i*.*e*., functions of the form

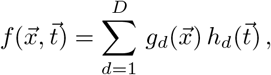

With 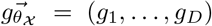 ϵ ℋ_χ_, 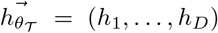ϵ ℋ_𝒯_ and D the common dimensionality of the two embeddings.

A second subspace ℋ_MLP_ corresponds to the MLP strategy of the two-branch architecture, in which the MLP is comprised of one hidden layer of size *L* ∈ ℕ, *i*.*e*., functions of the form

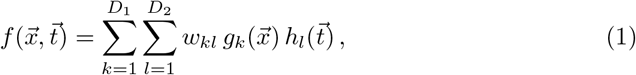

Where 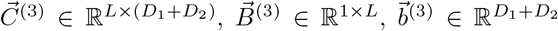, and 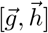 denotes vector concatenation of the D_1_-dimensional vector 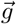 and the D_2_-dimensional vector 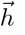. σ ○ represents an elementwise nonlinear transformation.

We introduce a third and final subspace ℌ _TP_ that defines the tensor product strategy of the two-branch architecture, in which compound and protein embeddings are aggregated by means of a tensor product, followed by a linear layer with a single output neuron. The resulting functions are of the form

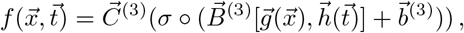

where 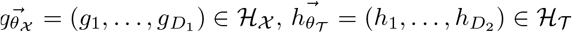, and 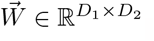.

### 2.2 Datasets

The proposed variants of the two-branch neural network architecture were evaluated on two benchmarks, Davis [32] and KIBA [33], because of their use as benchmarks in many of the studies presented in Table 1 and their varying numbers of compounds, proteins, and recorded affinities (see Table 2). The models were trained on the regression task, which aims to predict the affinity scores for compound-target pairs.

**Table 2:** Dataset statistics.

| Dataset | #drugs | #targets | #interactions |
| --- | --- | --- | --- |
| DAVIS | 68 | 379 | 30056 |
| KIBA | 2068 | 229 | 118254 |

### 2.3 Prediction settings

Collaborative filtering methods usually predict missing interactions between collections of users and items that have been observed during training (random-split), something that does not require any explicit feature representations. In contrast, the availability of such features in the typical DTI prediction task makes three additional prediction settings feasible [17]. The model could be expected to generate predictions for novel drugs (cold-drug) or novel targets (cold-target) that have not been observed during training. A fourth option that combines the strategies of the previous two is concerned with the prediction for pairs of novel drugs and targets that have not been observed during training. In the collaborative filtering area, these types of prediction settings are used less frequently but also witness a growing interest (cold start collaborative filtering). In this paper, we run experiments for the first three prediction settings: random-split, cold-drug and cold-target. The fourth prediction setting (combination of cold-drug and cold-target) was excluded from our analysis, as it is not implemented in the DeepPurpose library, which is used as the building block for protein and compound branches — see next paragraph. For each prediction setting, every dataset is split into training, validation and test sets (70-10-20%). However, the way the data is separated differs. For the cold-target setting, 70% of the targets only appear in the training dataset, 10% only appear in the validation datasets, and the remaining 20% only appear in the test set. In the cold-drug setting, the same ratios are used to split the drugs. Finally, for the random-split, the ratios are used to split the{*compound, protein, activity*} triplets in the dataset.

### 2.4 Benchmarking experiments

For the implementation of the two-branch architectures, the DeepPurpose library [15] was chosen as the starting point. A forked version of the repository, which has been heavily modified is available online^1^.

Since multiple branch pairings were possible, we included three combinations, reflecting varying degrees of descriptor and branch baseline complexity. These combinations included:

- An MLP compound branch on Morgan fingerprints [34] and an MLP protein branch on amino acid composition descriptors [35].
- A 1D Convolutional Neural Network (CNN) [36] compound branch on SMILES strings and a CNN protein branch on amino acid sequences.
- A Message-Passing Neural Network (MPNN) [37] compound branch on molecular graphs and a CNN protein branch on amino acid sequences.

### 2.5 Implicit feature representations

All the experiments mentioned above utilize different forms of explicit feature representations for both compounds and proteins, but not the structure of the interaction matrix. To better investigate the quality of these sources of information, we conducted additional experiments with the following differences:

- An MLP branch for the compounds and proteins that, instead of using explicit features, utilizes one-hot encoded dummy vectors. Since no generalization to new compounds or new proteins is possible when using this type of feature, we only focus on the random-split setting. When the dot product is used as the aggregation strategy the resulting architecture is a close analogue to traditional matrix factorization methods.
- A two-branch architecture where each branch is comprised of an internal two-branch model (Fig. 1.B). The internal model is designed to utilize the explicit and implicit features of the compounds/proteins, something that could potentially lead to improved performance. The MLP strategy is always used for the aggregation of the internal embeddings, while all three strategies of interest are available for the external embeddings.

### 2.6 Hyperparameter optimization

For the comparison of the three aggregation strategies across two DTI prediction datasets and the three prediction settings, we utilized random search as the hyperparameter optimization method of choice. For every optimization round, a budget of 100 configurations was allocated, with each experiment training for up to 100 epochs (early stopping on the validation loss was also used).

The hyperparameter ranges of every experiment had to be adapted based on the aggregation strategy and branch architectures used. The full details can be found in the Appendix. Every model was trained on a single GPU (either NVIDIA Ampere A100 or NVIDIA Volta V100), and all the results were logged using the Weights and Biases platform [38].

### 2.7 Metrics

To guarantee a consistent comparison across all our experiments, we utilized the same regression metrics used in the majority of the work presented in Table 1. These include the mean-squared error (MSE) and r-squared (R2) metrics.

## 3 Results

### 3.1 Universal approximation

Using existing mathematical results from [39, 40] as building blocks, one can easily show that all three aggregation strategies are universal approximators. We provide further details and formal derivations in Appendix, but summarize the main insights here. Broadly speaking, universal approximation theorems imply that neural networks can represent a wide variety of interesting functions when given appropriate weights. On the other hand, they typically do not provide a construction for the weights, but merely state that such a construction is possible.

In the setting of DTI prediction, universal approximation can only be guaranteed if the protein branch, compound branch and the aggregation strategy are universal approximators. For the formal results presented in the Appendix, we assume that the protein and compound branches are universal approximators, and we show that this is sufficient to prove universal approximation of the three aggregation strategies. For simplicity, we also assume that protein and compound feature vectors can be represented in Euclidean spaces, with multi-layer perceptrons as protein and compound branches. Similar universal approximation theorems could be easily derived for different activation functions [41, 42], non-Euclidean spaces [43], and other types of neural network architectures, such as deep convolutional neural networks [44].

### 3.2 Benchmarking experiments

In this section, we present extensive comparisons of the three embedding aggregation strategies on popular benchmark datasets from the field of DTI prediction. The experiments span two different DTI prediction datasets, three prediction settings (random, cold-drug, cold-target), as well as three different combinations of input feature representations and compound-protein branch architectures. Table 3 offers a quick summary of the results that have been obtained.

**Table 3:**
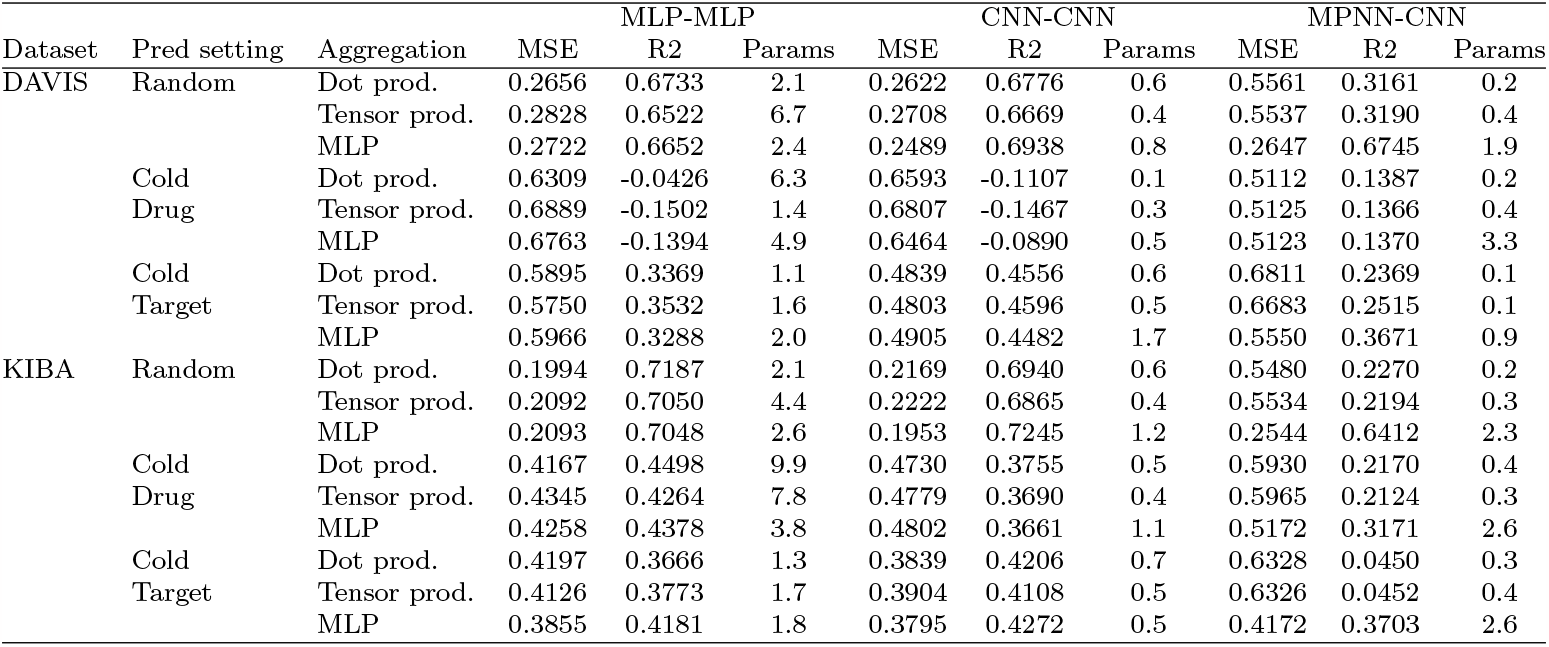
Performance results obtained for every possible combination of two datasets (KIBA, DAVIS), three prediction settings (random, cold-drug, cold-target), three embedding aggregation strategies and three compound-target branch pairs (MLP-MLP, CNN-CNN, MPNN-CNN). For every combination, the average MSE, R2 and number of trainable parameters of the top-5 performing (lowest overall loss) configurations are reported.

In the majority of cases and for both recorded metrics, we see that the dot product and tensor product strategies can be seen as competitive alternatives to the MLP. In many cases, they achieve superior performance. At the same time, none of the three strategies can be highlighted as the strategy of choice purely based on the final performance, as none of them consistently outperforms the others. These results confirm our theoretical findings presented in the Appendix, as all three strategies can approximate any target function. Interestingly, specific combinations of dataset, prediction setting and branch pair exist, in which the three stategies result in unexpected performance differences. More specifically, when we used an MPNN as the compound branch and a CNN as the protein branch, the dot product and tensor product strategies clearly failed to reach the performance of the MLP strategy in both randomly split datasets.

A more detailed investigation of the reasons behind this result as well as a potential remedy for the dot product and tensor product strategies are presented in a later subsection.

An important characteristic of any model that can influence many practical aspects of the training process is its overall capacity. Even though capacity measures of neural networks exist (e.g. Vapnik—Chervonenkis dimension [45]), they are primarily dealing with simple MLP architectures. Since the two-branch architecture we utilize can be equipped with more complex branches (CNNs, MPNNs, etc.) we decided to simplify things and instead use the total number of trainable parameters as the measure of model capacity. Since a smaller network with fewer parameters can result in reduced memory and lower computational requirements, a smaller model that can still achieve a similar performance compared with a larger counterpart is highly desirable. Our initial hypothesis was that the overhead of the MLP strategy introduced by the fullyconnected layers after the concatenation of the compound and protein embeddings would result in larger models. Based on the results shown in Table 3 the aforementioned competitiveness of the dot product strategy is usually achieved by small models. By accounting for this extra information, we can more confidently suggest the dot product strategy as a replacement for the MLP-based architecture.

Revisiting the close connection with recommender systems, a highly-cited publication by He et al. [18] first presented the dot product strategy as the simplest neural network approximation of matrix factorization. He et al. [18] then suggested the MLP strategy as a more powerful approach with the capacity to model more complex relationships between the items and users. This experimentally-backed strategy was then adopted by a series of subsequent publications [20–23] in the area of recommender systems, while proposals with the dot product strategy continued to be considered [24–26].

The superiority of the MLP aggregation strategy in the area of collaborative filtering was questioned by several subsequent publications. Rendle et al. [27] showed that, with careful hyperparameter selection, the dot product strategy could outperform the MLP strategy. They also pointed out that an MLP cannot trivially approximate the seemingly basic dot product operation. Xu et al. [28] offered a more rigorous comparison by investigating the limiting expressivity of each strategy, the convergence under the practical gradient descent algorithm, and the generalization potential. The two aforementioned publications approach the comparison of strategies exclusively in the area of recommender systems using benchmark datasets that are missing any explicit features for the users or items. In our investigation, which includes explicit feature representation and multiple generalization settings, we formulate similar conclusions as [27] and [28].

### 3.3 Implicit feature representations

Furthermore, Table 4 contains the results for two additional neural network configurations: two-branch neural networks that only use implicit features, and two-branch neural networks that use implicit and explicit features. So, in combination with the results of Table 3, which summarized the results for two-branch neural networks that only included explicit features, we compare here three types of two-branch neural networks. Overall, the initial setup with only explicit features gives the best results, but the differences between the three variants is small. The negligible differences let us conclude that adding implicit features does not have benefits for the datasets and models that we considered. Hovever, the neural network that only uses implicit features still yields a satisfactory performance, so a clear structure must be present in the interaction matrices of the two datasets.

**Table 4:** Performance results obtained for testing combinations of implicit and explicit compound and protein feature representations. The first combination uses to MLP branches that process dummy one-hot encoded features for the compounds and proteins. The second and third combinations represent two-level two-branch architectures. The outer level is comprised of the classic two-branch architecture with embeddings that can be combined using the three strategies we investigate. The two outer branches are composed of two-branch models that utilize both explicit and implicit features. For every combination, the average MSE, R2, and number of trainable parameters of the top-5 performing (lowest overall loss) configurations are reported.

| Dataset | Aggregation | MLP-MLP (implicit) |  |  | MLP-MLP (implicit+explicit) |  |  | CNN-CNN (implicit+explicit) |  |  |
| --- | --- | --- | --- | --- | --- | --- | --- | --- | --- | --- |
|  |  | MSE | R2 | Params (M) | MSE | R2 | Params (M) | MSE | R2 | Params (M) |
| DAVIS | Dot prod. | 0.2804 | 0.6551 | 5.2 | 0.2859 | 0.6484 | 5.2 | 0.2690 | 0.6692 | 2.3 |
|  | Tensor prod. | 0.3008 | 0.6301 | 9.6 | 0.3138 | 0.6141 | 9.6 | 0.3394 | 0.5825 | 1.3 |
|  | MLP | 0.3104 | 0.6182 | 8.3 | 0.3217 | 0.6043 | 8.3 | 0.3135 | 0.6145 | 2.0 |
| KIBA | Dot prod. | 0.2481 | 0.6500 | 7.2 | 0.2277 | 0.6789 | 7.2 | 0.2296 | 0.6761 | 3.5 |
|  | Tensor prod. | 0.2625 | 0.6297 | 12.9 | 0.2555 | 0.6396 | 12.9 | 0.2630 | 0.6290 | 2.6 |
|  | MLP | 0.2440 | 0.6559 | 12.0 | 0.2590 | 0.6347 | 12.0 | 0.2489 | 0.6490 | 3.2 |

In the last decade, matrix factorization methods have been extensively used to exploit the structure in the interaction matrix by decomposing it into two small matrices that contain the implicit features [46]. Formally speaking, the structure of the interaction matrix can be summarized using the singular values of that matrix. If most singular values differ from zero, the interaction matrix has a high rank, so approximating it as a product of two smaller matrices will lead to little predictive power and meaningless implicit features. Conversely, if most singular values are equal to zero, the interaction matrix has a low rank, and low-rank matrix approximation via matrix factorization will result in a good predictive performance and meaningful inplicit features.

Matrix factorization methods share major similarities with the two-branch architectures we consider in this work. The simplest version of the two-branch architecture, which uses only the implicit features and aggregates the embeddings via the dot product strategy, can be seen as a way of performing matrix factorization [18, 28, 47–49]. For explicit feature representations and other aggregation strategies, the link with matrix factorization is less obvious, and the models become more difficult to analyze in a formal way. However, we believe that the structure of the interaction matrix is also exploited by the models in that case, because low-dimensional embeddings of proteins and compounds are constructed. To our opinion, that’s the main reason why adding implicit features does not lead to performance gains in our experiments.

Let us remark that matrix factorization methods have been extensively used for DTI prediction. Early work by Cobanoglu et al. [50] used probabilistic matrix factorization combined with active learning without relying on compound or protein similarities. Ezzat et al. [51] proposed a graph regularized matrix factorization methods (and a weighted variant) to perform manifold learning and improving the performance in the cold-drug and cold-target prediction settings. More recent work by Mazzone et al. [52] used the NXTfusion [53] library that extends traditional matrix factorization methods in a non-linear fashion by inferring over an arbitrary number of data matrices, which are built as an entity relation graph. The data fusion step was performed by training a multi-task neural network that includes side information.

The concept of combining implicit and explicit information to improve the performance of DTI prediction tasks has also been incorporated into various kernel-based methods. Gonen [29] proposed a Bayesian formulation that combines kenrel-based nonlinear dimensionality reduction, matrix factorization and binary classification to predict interactions using only the chemical similarity between compounds and genomic similarity between proteins. Another kernel method, called NMTF-DTI [54], used a nonnegative tri-factorization technique based on Laplacian regularization and multiple similarity matrices for drugs and targets. A third approach defined the Gaussian interaction profile kernel [55] to capture topological information from the drug-target interaction network, and a variation of that method improved the predictive performance even further by incorporating extra sources of chemical and genomic information with additional kernels. These results do not agree with our findings, but the datasets used in most kernel-based papers were significantly smaller than those we analyzed.

Similar to the two-branch neural networks we discuss in this work, kernel methods enable the projection of the compounds and proteins into a shared space from which the predictions are generated. However, in two-branch neural networks, the branches “learn” the compound and protein embeddings, while kernel-based methods pre-compute these embeddings by specifying a specific kernel. In general one can assume that learning the embeddings is better than defining them based on a kernel, especially when datasets are large. That’s probably the reason why deep learning methods have won the battle against kernel methods in DTI prediction. Furthermore, it is worth mentioning that the use of multiple kernels for drugs and targets with the goal of improving performance [56, 57] shows many similarities with multi-modal neural network architectures that utilize multiple branches (one per feature representation) before the aggregation step [58–61]. Both concepts essentially try to boost performance by including different types of information from the same entity.

### 3.4 Embedding visualizations

To better understand the similarities and differences between the MLP and dot product, we experimented with various visualizations and included the most interesting examples in this work. From this point onwards, our analysis skips the tensor product, as it does not provide any performance gains or distinct characteristics compared with the MLP and dot product strategies. In their simplified form, one of the key conceptual distinctions among these strategies lies in the location of the learning process. In the dot product strategy, this is exclusively done in the two branches since the aggregation strategy is just the dot product operation. In contrast, the MLP strategy shares the learning between the two branches and the fully-connected layers after the concatenation of the embeddings. A basic visualization of the compound-protein embeddings obtained from the fully-connected layers (Fig. 2) of a well-performing MLP strategy shows an improving separation of the active and inactive compound-protein pairs the closer we get to the output node.

**Fig. 2:**
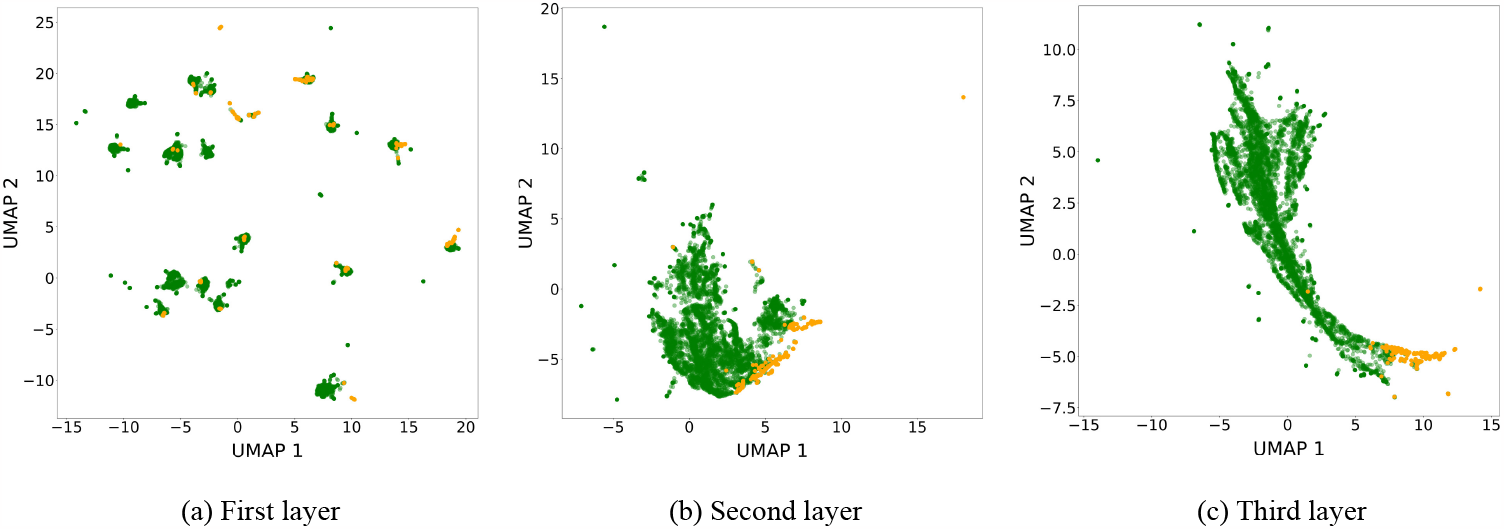
UMAP visualization of the test compound-protein embeddings extracted from the fully-connected layers of the best MLP configuration trained on the DAVIS dataset. The configuration uses CNN compound and protein branches. The points are colored using the binarized affinities (orange for active and green for inactive). From the figure, we observe that starting from the first layer (after the concatenation of the compound and protein branch embeddings) to the third, the separation of the two classes becomes clearer.

To investigate the quality of the compound embeddings, we experimented with another type of visualization in Figure 3. The four subplots visualize the affinity scores of four proteins (with the highest number of recorded affinities in the KIBA dataset) after the Uniform Manifold Approximation and Projection (UMAP) [62] is applied on the compound embeddings.

**Fig. 3:**
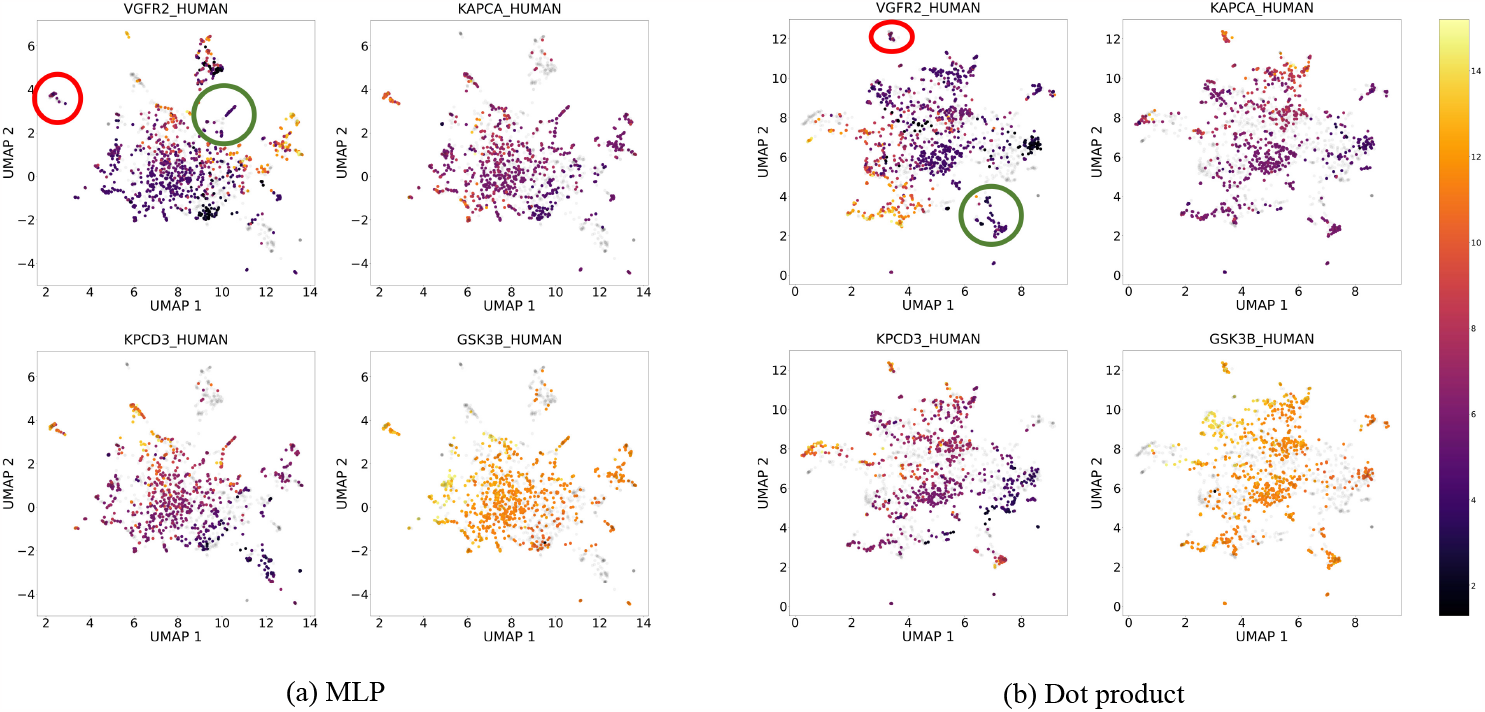
UMAP visualization of the compound embeddings extracted from the last fully-connected layer of the compound branch (CNN) of the best MLP (left) and dot product (right) configurations trained on the KIBA dataset. The points are colored based on the affinities of the four proteins in the training set with the highest number of recorded entries. Comparing the two strategies, we observe that both create clusters of similar quality.

Even though the affinity differences observed in compound clusters between the four proteins are quite interesting, we focus on the relative comparison of the learned clusters. Based on the aforementioned discussion about the location of the learning process, the well-defined clusters of the compound embeddings from the dot product strategy are largely expected, as the two branches are exclusively responsible for learning to predict affinities. At the same time and, somewhat surprisingly, the visual inspection of the compound embeddings obtained from the MLP strategy shows similar embedding quality as the dot product, even though the fully-connected layers after the embedding concatenation share the learning responsibility.

### 3.5 Oversmoothing effect and its importance when selecting an embedding aggregation strategy

For specific combinations of dataset, prediction setting and branch pair, Table 3 includes dot product and tensor product examples that are clearly outperformed by the MLP aggregation strategy. These cases include the use of the MPNN compound branch and the CNN protein branch on the randomly split version of the two datasets. As the CNN protein branch had been successfully used in the CNN-CNN combination, we focused our efforts on the MPNN branch as the defective model. In an effort to increase its capacity, we first tested configurations with deeper MPNNs, something that ultimately did not yield any improvements. Our hypothesis was that the oversmoothing effect that many graph neural networks suffer from [63] made any attempt to increase the capacity fail performance-wise. This is shown in the two plots of Figure 4, where points from models that use the dot product strategy and varying sizes of the MPNN branch architecture fail to approach the performance the MLP strategy achieves. Assuming that the superior performance of the MLP-based architecture was a result of the fully-connected layers, we decided to test new dot-product-based configurations by appending fully-connected layers after a smaller MPNN model and before the embedding aggregation step. With this strategy, we managed to both increase the capacity of the compound branch and avoid the over-smoothing effect that larger MPNN models suffer from. Referring back to Figure 4, these modified architectures colored in red show major improvements, as they became comparable performance-wise with the MLP strategy.

**Fig. 4:**
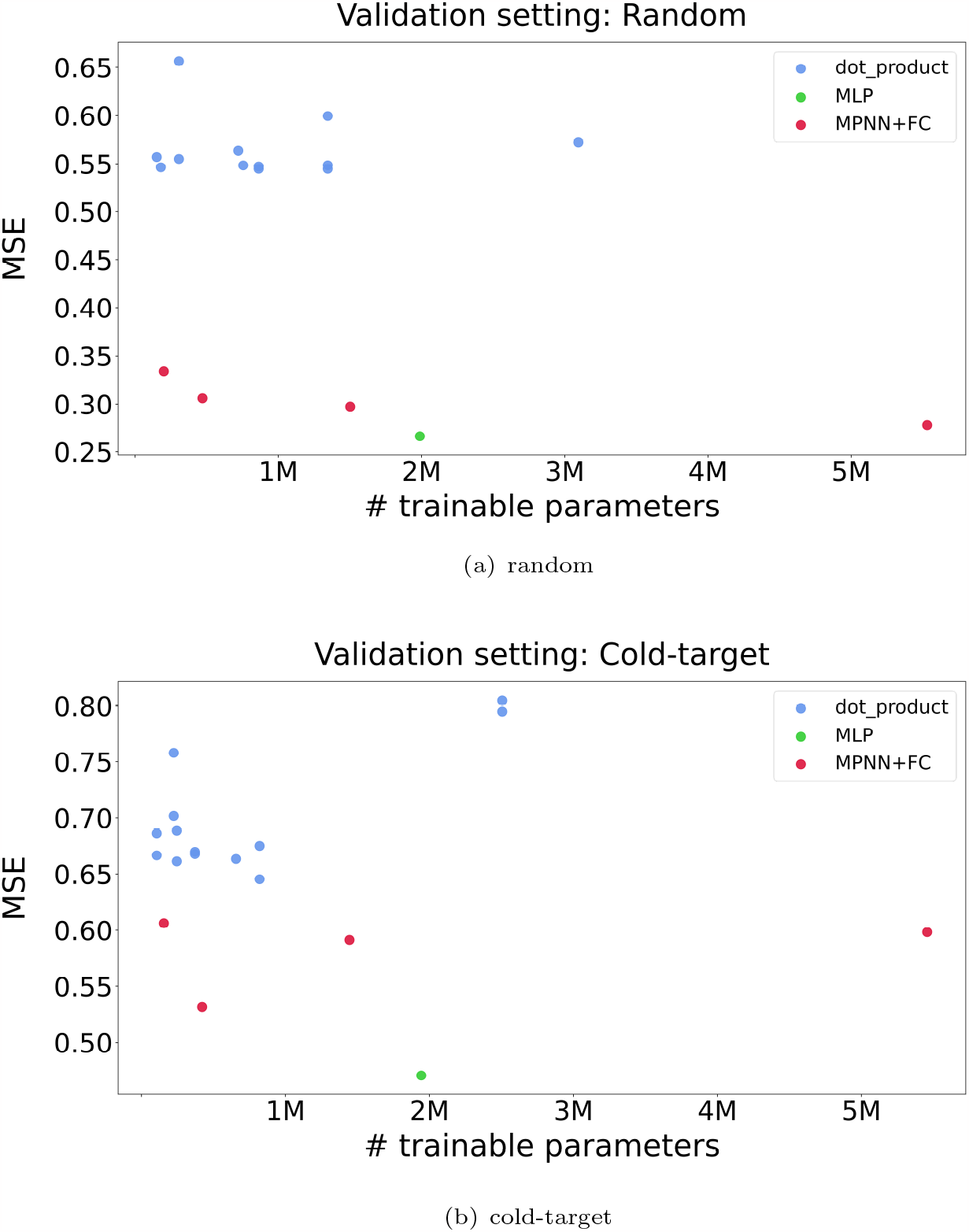
Plot of the capacity of different models against their test MSE. All the points represent experiments with an MPNN as the compound branch and a CNN as the protein branch. The point colored green represents a well-performing experiment of the MLP strategy, while the blue points represent dot product strategies with similar branch configurations. The results illustrate the inability of the dot product to achieve comparable performance with the MLP strategy when one of the branches fails to utilize its input properly. Even after testing dot-product-based configurations with varying capacity (increasing the MPNN depth) exceeding that of the MLP benchmark, the performance does not improve. Our hypothesis is that the main reason for the underperforming experiments that use the dot product is the over-smoothing effect that graph neural networks suffer from. This assumption is supported by the significant improvements that a small modification attains (colored red). Instead of increasing the capacity of the MPNN directly, we instead append fully-connected layers immediately after it and before the embedding aggregation operation, thus increasing the overall capacity of the compound branch and bypassing the MPNN. The experiments colored red demonstrate that the dot product strategy can reach a comparable performance as the MLP strategy while keeping the overall capacity low.

The underlying message of this exploration is that the dot product has the inability to counterbalance weaknesses that may arise from the branches, as it lacks the trainable parameters that the MLP strategy has. This problem can be tackled by increasing the capacity of the defective branch. However, cases still exist where this strategy is inadequate because of branch-specific weaknesses (e.g., over-smoothing in GNNs). In such cases, it is suggested to increase the capacity of the defective branch with fully-connected layers, replace the branch altogether.

## 4 Conclusion

In this work, we analyzed alternatives to the traditional embedding aggregation strategy that has dominated the two-branch architectures in the field of DTI prediction. We presented formal and experimental results which show that all three analyzed strategies can be used to aggregate embeddings. We identified conditions under which a particular type of aggregation strategy might outperform others, and we presented various visualizations. We believe that this work can be the first step in convincing the DTI prediction community to also focus on the embedding aggregation options. Even though this may not seem vital when aggregating only two embeddings, it can become an important choice in multi-modal architectures or when more than two embeddings have to be aggregated (e.g. drug-drug-protein interaction prediction).

With regard to future work, we intend to test attention-based embedding aggregation methods, which have become popular in other application domains. Furthermore, increasing the number of embeddings that are aggregated is also an interesting avenue we intend to explore. This approach is used when different representations are available for the same entity. For example, different feature representations for a given compound (Morgan fingerprint, 2D image, molecular graph) could be combined in different ways. Thus, the order and strategy used at every stage of aggregating embeddings is a complex task. Finally, the evaluation of the added value that pre-trained embeddings from different strategies can bring to transfer learning tasks is an interesting topic that we intend to investigate in future work.

## Declarations

- Ethics approval and consent to participate: Not applicable
- Consent for publication: Not applicable.
- Availability of data and materials: The KIBA and DAVIS datasets are publicly available https://github.com/futianfan/DeepPurpose Data. The scripts used to perform the experiments are also available online https://github.com/diliadis/DeepPurpose as a forked and modified repository of the original Deep Purpose project.
- Competing interests: The authors declare that they have no competing interests.
- Funding: This research received funding from the Flemish Government under the

“Onderzoeksprogramma Artificiele Intelligentie (AI) Vlaanderen” programme.

1. Authors’ contributions: D.I. performed the research. D.I., W.W, T.P. and B.B. contributed to the manuscript, read and approved the final version.

## Footnotes

1 https://github.com/diliadis/DeepPurpose

